# N^1^-methylpseudouridine-incorporated mRNA enhances exogenous protein expression and suppresses immunogenicity in primary human fibroblast-like synoviocytes

**DOI:** 10.1101/2022.03.22.485393

**Authors:** Sho Mokuda, Hirofumi Watanabe, Hiroki Kohno, Michinori Ishitoku, Kei Araki, Shintaro Hirata, Eiji Sugiyama

## Abstract

Studies conducted using murine arthritis models have indicated that the use of *in vitro*-transcribed messenger RNA (IVT mRNA) is an effective therapeutic approach for joint diseases. However, the use of IVT mRNA in human synovial cells has not been widely studied. Recently, the outbreak of the novel coronavirus disease has accelerated the development of innovative mRNA vaccines such as those containing a modified nucleic acid, N^1^-methylpseudouridine-5′-triphosphate (m1ψ). IVT mRNA is an attractive tool for biological experiments and drug discovery. To verify the protein expression of IVT mRNA *in vitro*, primary cultured human fibroblast-like synoviocytes (FLS) were transfected with enhanced green fluorescent protein (EGFP) mRNA with or without m1ψ incorporation. EGFP was detected using western blotting and fluorescence microscopy. A multiplex assay was performed to comprehensively understand IVT mRNA-induced immunogenicity. FLS transfected EGFP mRNA containing m1ψ generated higher levels of EGFP than unmodified EGFP mRNA or control RNAs. The multiplex assay of the FLS culture supernatant revealed that concentrations of IL-6, TNF-α, and CXCL10 were upregulated by unmodified EGFP mRNA, whereas they were suppressed by EGFP mRNA with m1ψ. Overall, m1ψ incorporation enhanced protein expression and decreased cytokine expressions in primary cultured FLS. The findings may contribute to arthritis research.

## Introduction

Studies conducted using murine arthritis models have indicated that the use of *in vitro*-transcribed messenger RNA (IVT mRNA) is an effective therapeutic approach for joint diseases ^1,2^. IVT mRNA is synthesized *in vitro* from template DNA. When IVT mRNA penetrates the cell membrane and enters the cytoplasm, it instantly initiates the production of foreign proteins using intracellular ribosomes. Unlike plasmid DNA and viral vectors, IVT mRNA can function without entering the nucleus and is degraded through the endogenous metabolic pathway after protein production ^3^. These therapeutic strategies using IVT mRNA are attractive and have been examined *in vivo*. However, the *in vitro*-use of IVT mRNA in human synoviocytes has been poorly studied ^4^.

IVT mRNA introduced into the cytoplasm stimulates intracellular innate immune responses through pattern-recognition receptors (PRRs) ^3^. Exogenous single-stranded RNA (ssRNA) is known to react with Toll-like receptor 7 (TLR7), TLR8, protein kinase R (PKR) and retinoic acid-inducible gene-I (RIG-I) ^5–10^. In addition, when IVT mRNA forms a hairpin or partially forms double-stranded RNA (dsRNA), it may also be recognized by PRRs such as TLR3, RIG-I, PKR, 2′–5′–oligoadenylate synthetase (OAS)/ribonuclease L and melanoma differentiation-associated protein 5 (MDA5) ^11–14^. It has been reported that the incorporation of modified nucleic acids, such as pseudouridin-5′-triphosphate (ψ), into IVT mRNA is an effective method to reduce immunogenicity ^10,15,16^. The advantages of ψ and 5-methylcytidine-5′-triphosphate have also been reported ^1,17^.

Recently, the novel coronavirus disease 2019 pandemic, which has been ongoing since December 2019, has led to the development of innovative mRNA vaccines such as BNT162b2 and mRNA-1273, containing N^1^-methylpseudouridine-5′-triphosphate (m1ψ) ^18,19^. These vaccines seldom have only a few negative effects on the human body, at least in the short term (within approximately 1 year), indicating the safety of m1ψ. The incorporation of m1ψ into IVT mRNA has been highlighted as a useful method for enhancing protein expression ^20–27^; m1ψ has a unique benefit for therapeutic methods using IVT mRNA.

In this study, we aimed to evaluate the efficiency of exogenous gene expression and immunogenicity of m1ψ-incorporated IVT mRNA using primary cultured human fibroblast-like synoviocytes (FLS). Our study demonstrated that lipofection of m1ψ-incorporated IVT mRNA is a potential *in vitro* gene transfer technique for FLS.

## Materials and Methods

### Ethics

This study was approved by the clinical ethics committee of Hiroshima University Hospital (approval number: E-668; approval date: February 1, 2017). The methods were performed in accordance with the approved guidelines. We collected synovial tissues from patients with rheumatoid arthritis (RA) who fulfilled the classification criteria of the American College of Rheumatology 1987 ^28^. Synovial tissues were harvested from patients with RA who underwent total joint replacement or synovectomy after they provided informed consent and signed a written consent form.

### Preparation of primary cultured FLS

For FLS preparation, synovial tissues from three patients with RA were minced and incubated with 1 mg/mL collagenase/dispase (Roche, Indianapolis, IN, USA) in phosphate-buffered saline (pH7.2, FUJIFILM Wako Pure Chemical Co.) for 1 h at 37°C. The synovial cells were then filtered, washed, and cultured. During incubation, the supernatant was replaced frequently to withdraw any nonadherent cells. Adherent FLS were then subcultured at 1:3 after reaching 80% confluence and passaged. FLS were cultured in Dulbecco’s modified Eagle medium (DMEM; FUJIFILM Wako Pure Chemical Co.) supplemented with 10% fetal bovine serum (Sigma-Aldrich) and penicillin/streptomycin (FUJIFILM Wako Pure Chemical Co.). FLS were used in the experiments at passages 3–4 and starved in DMEM containing 0.5% fetal bovine serum for 24 h before measuring mRNA and protein expression. Both cells were incubated at 37 °C under 5% CO_2_.

### Preparation of IVT mRNA and transfection into the cultured cells

To prepare templates for IVT mRNA with a cap structure and poly A sequence, the following plasmids were previously constructed in our laboratory: pcDNA3-A124 and pcDNA3-enhanced green fluorescent protein (EGFP)-A124 vectors, containing both untranslated regions and poly A contiguous sequences ^1,4,29^. Template plasmids were linearized, and *in vitro* transcription was performed using the HiScribe T7 ARCA mRNA Kit (New England Biolabs, Ipswich, MA, USA) to generate IVT mRNA according to the manufacturer’s instructions. m1ψ was purchased from TriLink BioTechnologies (San Diego, CA, USA). Components of the mixture except for the template plasmids, reaction buffer, and T7 polymerase were as follows. (1) Unmodified IVT mRNA: 4 mM cap analog (anti-reverse cap analog; 3′-O-Me-m7G(5′)ppp(5′)G), 1.25 mM ATP, 1 mM GTP, 1.25 mM CTP, and 1.25 mM UTP. (2) The IVT mRNA (m1ψ): 4 mM cap analog (anti-reverse cap analog), 1.25 mM ATP, 1 mM GTP, 1.25 mM CTP, 1.25 mM UTP, and 2.5 mM m1ψ. After *in vitro* transcription at 37 °C for 1 h, IVT mRNA was purified using a Monarch RNA Cleanup Kit (New England Biolabs). IVT mRNA was then transfected into cells using Lipofectamine MessengerMAX (Thermo Fisher Scientific, Waltham, MA, USA) according to the manufacturer’s instructions. Briefly, mRNA was mixed with Lipofectamine reagent at a ratio of 1:3 (pmol mRNA: µL Lipofectamine) in serum-free DMEM. The complex was allowed to form for 20 min at 25 °C. At 24 h after transfection, both cells and their culture supernatants were harvested for the subsequent experiments.

### Western blotting

The cells were plated and washed with phosphate-buffered saline before collection. Proteins from the cultured cells were processed using SuperSep Ace 15% precast gel (FUJIFILM Wako Pure Chemical Co.) and transferred on to a polyvinylidene fluoride membrane. The membranes were probed with anti-GFP (0.5 µg/mL, chicken polyclonal; Genscript, Piscataway, NJ, USA) and anti-β-actin (1 µg/mL, mouse monoclonal, clone. AC-15; Sigma-Aldrich) antibodies. Horseradish peroxidase-conjugated secondary antibodies (Jackson ImmunoResearch, West Grove, PA, USA) were then added. Horseradish peroxidase activity was detected using ECL prime reagents (Cytiva, Tokyo, Japan), followed by imaging with Image Quant LAS 500 (Cytiva).

### Measurement of EGFP fluorescence intensity

To observe the fluorescence of live cells, the culture medium was replaced with FluoroBrite DMEM (Thermo Fisher Scientific), and then imaged under the green channel of a ZOE Fluorescent Cell Imager (Bio-Rad Laboratories, Hercules, CA, USA).

### Measurement of cell viability

To measure the viability of the cultured cells, the transfected cells plated in 96-well black plates were examined using a CellTiter-Blue Cell Viability Assay kit (Promega, Madison, WI, USA). Briefly, 20 μL of CellTiter-Blue Reagent (resazurin) was added to 100 μL of the medium in each well and incubated at 37 °C for 1 h. Resazurin is reduced to resorufin by the aerobic respiration of metabolically active cells. The fluorescence of resorufin was measured using a SpectraMax iD3 system (Molecular Devices) at the excitation and emission wavelengths of 550 and 595 nm, respectively.

### Immunocytochemistry staining

FLS were cultured on cover slips placed in a six-well tissue culture plate. After transfection with IVT mRNA, the cells were incubated for 24 h. The cells were fixed with a mixture of acetone and methanol for 20 min at 25°C, and then blocked with 5% Blocking One P reagent (Nacalai Tesque, Kyoto, Japan) diluted with Tris-buffered saline for 1 h at 25 °C. The cells were incubated overnight at 4°C with the primary antibodies against GFP tag (8 µg/mL; rabbit polyclonal; Proteintech, Rosemont, IL, USA), and then with Alexa Fluor 647-conjugated anti-rabbit IgG antibodies (4 µg/mL; host: donkey; Abcam, Cambridge, UK) for 1 h at 25°C. The cells on the coverslips were mounted on glass slides with VECTASHIELD mounting medium containing 4′,6-diamidino-2-phenylindole (DAPI) (Vector, Burlingame, CA, USA). Anti-GFP antibody signals were detected using a digital microscope VHX-7000 (KEYENCE, Osaka, Japan).

### Multiplex assay by fluorescence-activated cell sorting (FACS)

The concentrations of interleukin-1β (IL-1β), IL-6, IL-8, IL-10, IL-12p70, interferon-α2 (IFN-α2), IFN-β, IFN-λ1, IFN-λ2/3, IFN-γ, tumor necrosis factor-α (TNF-α), C-X-C motif chemokine ligand 10 (CXCL10), and granulocyte macrophage colony-stimulating factor (GM-CSF) in the supernatant of cultured cells were measured using a LEGENDplex Multi-Analyte Flow Assay Kit, Human Anti-Virus Response Panel (Biolegend, San Diego, USA). The assay was performed in a V-bottom plate according to the manufacturer’s protocol. Data were acquired using a Cytoflex flow cytometer (Beckman Coulter, Brea, CA, USA) and analyzed using Legendplex Data Analysis V7.1 software (Biolegend). The sensitivity of the assay, indicated in parentheses, was as follows: IL-1β (19.3 pg/mL), IL-6 (11.7 pg/mL), IL-8 (5.7 pg/mL), IL-10 (15.4 pg/mL), IL-12p70 (3.7 pg/mL), IFN-α2 (2.9 pg/mL), IFN-β (105.7 pg/mL), IFN-λ1 (47.7 pg/mL), IFN-λ2/3 (270.5 pg/mL), IFN-γ (39.8 pg/mL), TNF-α (16.9 pg/mL), CXCL10 (16.3 pg/mL), and GM-CSF (15.6 pg/mL).

### Statistical analysis

All statistical analyses were performed using Student’s *t*-test. All graphs show the results of one representative experiment from several individual experiments. The results were analyzed and processed using GraphPad Prism 9 software (GraphPad, Inc., La Jolla, CA, USA).

## Results

### IVT mRNA induction in primary cultured FLS

We prepared the following four types of IVT mRNA: mock mRNA with or without m1ψ-incorporation and EGFP mRNA with or without m1ψ-incorporation. Mock mRNA, which cannot produce any protein, is approximately 400 nucleotides (nt) and EGFP mRNA is approximately 1,100 nt. We examined the expression level of EGFP in transfected primary cultured FLS. FLS were harvested from human synovial tissues, passaged, and transfected with IVT mRNA. Samples were collected at 24 h after transfection. As shown in **Fig 1a** (western blotting) and **Fig 1b** (fluorescence microscopy), EGFP expression in EGFP mRNA (m1ψ)-transfected FLS was higher than that in the negative control groups or unmodified EGFP mRNA-transfected FLS. Unexpectedly, FLS produced green auto-fluorescence in the control groups. Then, we also performed immunocytochemistry using Alexa Fluor 647 to calculate the transfection efficiency. The efficiency of EGFP mRNA (m1ψ)-transfected FLS was more than 80% (**Fig 1c**). And, the viability of these cells was more than 90% (**Fig 1d**).

**Fig 1.**
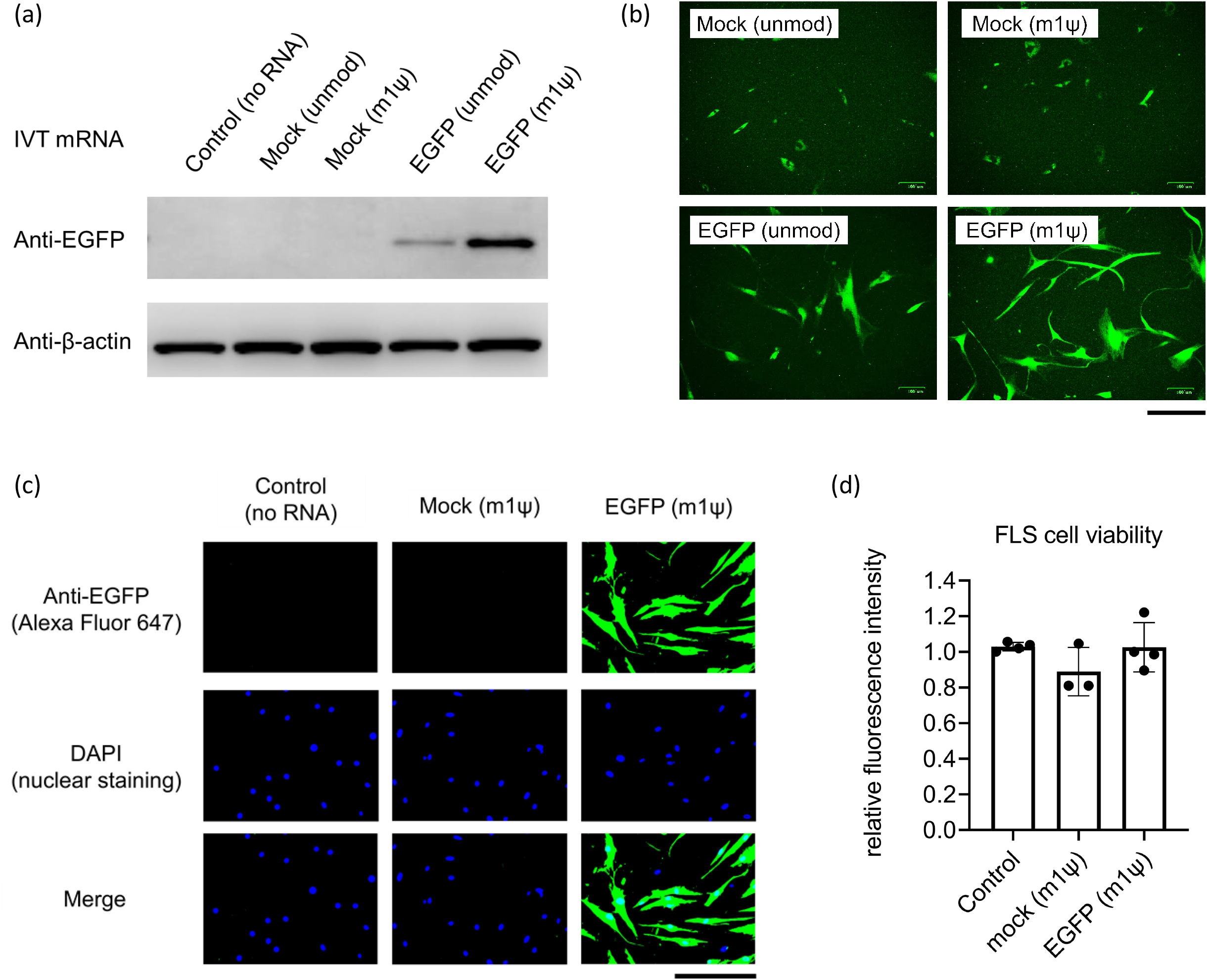
EGFP mRNA transfection into primary human fibroblast-like synoviocytes (FLS) FLS were transfected with *in vitro*-transcribed mRNA (IVT mRNA) and harvested at 24 h after transfection. (a) Western blotting of EGFP. (b) Images of FLS captured by fluorescence microscopy. Scale bar, 0.2 mm. (c) Immunocytochemical staining for EGFP. Scale bar, 0.2 mm. (d) Measurement of cell viability using the CellTiter-Blue Cell Viability Assay kit (n = 3–4, mean ± SEM).

These findings indicate that IVT mRNA containing m1ψ can induce adequate foreign protein expression with acceptable cell viability in primary cultured FLS.

### Comprehensive analysis of IVT mRNA-stimulated cytokine production in FLS

Introduced IVT mRNA is considered to be recognized by some PRRs and induce post-viral infection-like responses, which can activate two main cascades, namely, nuclear factor-kappa B (NF-κB)-mediated signaling and interferon-regulatory factor (IRF)-mediated signaling ^3,27^. To clarify these responses against IVT mRNA, we used a multiplex assay to measure 13 different proteins in the culture supernatants from 0 to 24 h after transfection: IL-1β, IL-6, IL-8, IL-10, IL-12p70, IFN-α2, IFN-β, IFN-λ1, IFN-λ2/3, IFN-γ, TNF-α, CXCL10, and GM-CSF. IL-6, TNF-α, CXCL10, and IL-8 were detected, whereas the other nine proteins were below the detection limits. Unmodified mock mRNA increased the protein levels of IL-6 and CXCL10. Moreover, unmodified EGFP mRNA increased the protein levels of IL-6, TNF-α, and CXCL10 (**Fig 2a–c**). Both mock and EGFP mRNAs incorporated with m1ψ downregulated this elevation. IL-8 showed high endogenous excretion in the no RNA control group, and it did not increase in the presence of unmodified IVT mRNA (**Fig 2d**).

**Fig 2.**
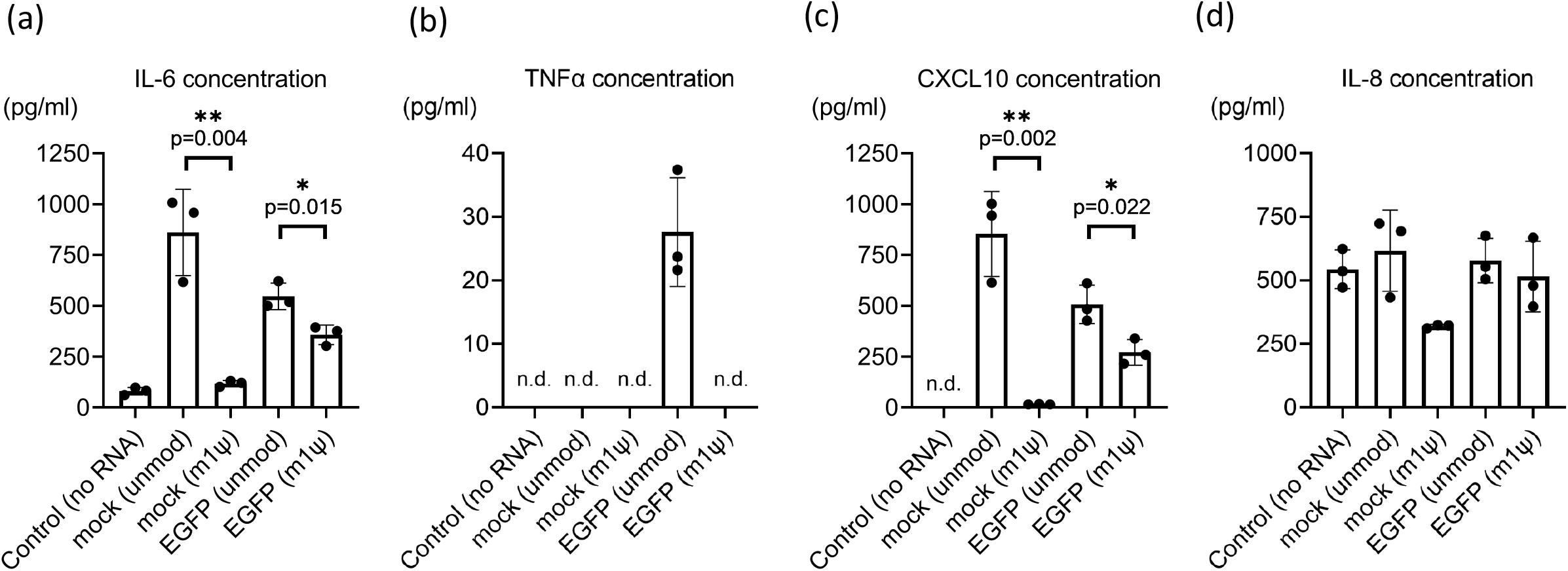
Multiplex assay for *in vitro*-transcribed mRNA (IVT mRNA)-transfected fibroblast-like synoviocytes (FLS) culture supernatant to measure cytokine concentrations. The cytokine concentrations in the FLS culture supernatant were measured using a LEGENDplex Multi-Analyte Flow Assay kit. The samples contained humoral factors for 24 h after transfection (n = 3). (a) IL-6 (pg/mL), (b) TNF-α (pg/mL), (c) CXCL10 (pg/mL), and (d) IL-8 (pg/mL) were detected. IL-1β, IFN-λ1, IL-12p70, IFN-α2, N-IFN-λ2/3, GM-CSF, IFN-β, IL-10, and IFN-γ were below the detectable limit. Data are presented as mean ± SEM. *t*-test, *p < 0,05, **p < 0.01.

In summary, unmodified IVT mRNA simultaneously elevated both NF-κB-mediated pro-inflammatory cytokines, IL-6 and TNF-α, and an IRF-mediated chemokine, CXCL10. In addition, m1ψ-incorporation into IVT mRNA suppressed this elevation.

## Discussion

From 1993 to 2000, IVT mRNA was developed as a vaccination approach against cancer and infectious diseases, mainly because it does not undergo mutagenesis and can transiently produce various exogenous protein ^30–35^. In contrast, IVT mRNA itself can induce unfavorable immunogenicity. When IVT mRNA is used as a vaccine, these immune-stimulatory responses may act as adjuvants and promote adaptive immune responses against the introduced exogenous protein ^36^. In contrast, when IVT mRNA is treated as a gene replacement therapy, additive immune responses are not necessary, and it may have disadvantages.

Inflammation exacerbates the pathophysiology of joint diseases, and human RA-derived FLS reportedly produce high levels of IL-6 and CXCL10 ^37–39^. Therefore, it is desirable that IVT mRNA has low immunogenicity when developing a therapeutic strategy involving gene replacement for arthritis. To verify the usefulness of m1ψ-incorporated IVT mRNA for cultured cells, we utilized primary cultured human FLS. To date, transfection of both unmodified IVT mRNA and plasmid DNA using lipofection in FLS is technically difficult, and these molecules must be introduced using electroporation or viral vectors. Our results support that m1ψ-modified IVT mRNA-lipofection method is convenient, facilitates high protein production, and as noted below, has low immunogenicity.

To avoid immune responses induced by IVT mRNA, it is necessary to block the interaction between these single-stranded RNAs, which partially form double strands, and PRRs such as TLR3, TLR7, TLR8, RIG-I, PKR, OAS/ribonuclease L, and MDA5 ^5,8,10,14,15,40^. One strategy is using modified nucleic acids, such as m1ψ. However, the biological functions of m1ψ-incorporated IVT mRNA in cultured human cells are not well understood. Most studies have indicated that this mRNA increases expression efficiency; however, its ability to suppress innate immune responses has not been evaluated in detail. Particularly, there are only a few comprehensive studies on cytokine production induced by IVT mRNA-stimulated cells at the protein level. Kormann et al. reported both elevated TNFα and IL-8 concentrations in human monocyte culture supernatant stimulated by IVT mRNA ^41^. In the present study, we analyzed the comprehensive immune response induced by IVT mRNA. IL-6, TNF-α, and CXCL10 were particularly noticeable among the 13 proteins tested. Measurement of the protein concentrations and mRNA expression revealed that m1ψ incorporation can repress the immunogenicity of IVT mRNA. The following findings of *in vitro* experiments have been reported: *IFN-β* and C-C motif chemokine ligand 5 (*CCL5*) expression is suppressed in IVT mRNA (m1ψ)-transfected human A549 cells; both *IFN-α* and *IFN-β* expression is downregulated in IVT mRNA (m1ψ)-transfected rat cardiomyocytes; and *CXCL10* expression is downregulated in IVT mRNA (m1ψ)-transfected human monocyte-derived macrophages *in vitro* ^20,24,26^. These results are consistent with our results.

In conclusion, we demonstrated that m1ψ incorporation enhances exogenous protein expression and suppresses immunogenicity in primary human cells. Thus, m1ψ-incorporated IVT mRNA will be useful for primary culture experiments, including arthritis research.

## Acknowledgments

We would like to thank Shigeru Miyaki (Hiroshima University) for the preparation of experimental devices.

## Statements and Declarations

### Availability of data and materials

The source data for most of the figures are available from the authors.

### Competing Interests

The authors declare no conflicts of interest associated with the manuscript.

### Authors’ contributions

SM, HW, HK, MI, and KA performed the experiments and analyzed the data. SM, HW, SH, and ES planned the experiments and wrote the manuscript.

### Funding

This research was funded by JSPS KAKENHI (Grant Number 19K18499 to S.M.), Mitsubishi Foundation to S.M., Takeda Science Foundation to S.M., Mochida Memorial Foundation for Medical and Pharmaceutical Research to S.M., Japanese Respiratory Foundation Grant to S.M., Japan Rheumatism Foundation to S.M., The Japan College of Rheumatology Grant for Promoting Research for Early RA to S.M., and The Nakatomi Foundation to S.M.

## Notes

### Competing Interest Statement

The authors have declared no competing interest.

